# The endemic plant species of Ebo Forest, Littoral Region, Cameroon with a new Critically Endangered cloud forest shrub, *Memecylon ebo* (Melastomataceae-Olisbeoideae)

**DOI:** 10.1101/2023.12.20.572583

**Authors:** Robert Douglas Stone, Barthelemy Tchiengué, Martin Cheek

## Abstract

We emphasise the urgent need to conserve the Ebo forest (Littoral Region, Cameroon), which holds 10 strict endemic plant species and 15 near endemics for a total of 25, a very high number far exceeding the threshold for its recent status as an Important Plant Area (IPA). We describe a further strict endemic species from the Ebo Forest, *Memecylon ebo* sp. nov. (Melastomataceae-Olisbeoideae) placed in sect. *Afzeliana* due to its ellipsoid blue-green fruits. The yellow petals and jade green anther-connectives of *M. ebo* are unique in the genus *Memecylon* as a whole, among its >400 species ranging overall from Africa to the western Pacific. *Memecylon ebo* is assessed as Critically Endangered (CR) using the 2012 IUCN standard due to the small range size and the extremely high and ongoing threats of logging at Ebo, and also due to subsequent threats of potential oil palm plantation and mining projects. With the addition of *Memecylon ebo*, the tally of Critically Endangered plant species recorded from Ebo forest is now the highest of any IPA in Cameroon, equalling that of, Ngovayang with 24 CR species.

## Introduction

The genus *Memecylon* L. (Melastomataceae-Olisbeoideae) is distributed throughout the Old World tropics, reaching 25°N latitude in NE India, N Bangladesh and S China, 32° S latitude in southern Africa and extending to the western Pacific in the Solomon Islands, Fiji and Tonga. The > 400 recognized species are shrubs or small trees mainly of surviving areas of tropical forest. Since January 2020, 66 names of *Memecylon* new to science have been published, mainly from Madagascar (Stone 2020, 2022a, 2022b, 2023) and also from India (IPNI, continuously updated). In accordance with morphological and molecular findings (Jacques-Félix 1978; Bremer 1982; Stone 2006, 2014a, 2022c; Stone & Andreasen 2010), *Memecylon* is now circumscribed to exclude the monospecific western and central African genus *Spathandra* Guill. & Perr., the paleotropical *Lijndenia* Zoll. & Moritzi, and the African-Madagascan *Warneckea* Gilg. The members of *Memecylon* sensu stricto are characterized by a combination of very hard wood; leaves opposite, estipulate, and apparently 1-nerved (less often “subtrinerved” sensu Jacques-Félix et al. 1978; Jacques-Félix 1983, 1985); a general lack of indumentum; flowers small and 4-merous; anther-connectives enlarged and with a dorsal oil-gland (or with gland reduced or absent in some species or species-groups); and fruits baccate with 1–few large seeds and embryo foliaceous and convoluted.

In continental Africa, *Memecylon* sensu stricto holds 73 species and is entirely sub-Saharan and mainly tropical, extending in the west from the Republic of Guinea with seven species (Gosline et al. 2023a, 2023b) and east to Tanzania with 15 species (POWO, continuously updated). *Memecylon* is entirely absent from NE Africa, including Ethiopia and Somalia and S. Sudan (e.g. Darbyshire et al. 2015). Taxon diversity in Africa is highest in Cameroon, with 32 species (Onana 2011). In contrast, for the moment only 21 species are listed for neighbouring Gabon (Sosef et al. 2006). Many of the Cameroon species are narrow endemics and highly threatened by forest habitat clearance. The Cameroon Plant Red Data book includes 12 *Memecylon* species, all but one of which is either Endangered or Critically Endangered (Onana & Cheek 2011).

Within Cameroon, the highest diversity of species of *Memecylon* is held within the Cross-Sanaga River Interval (Cheek et al. 2001) as is recorded also in numerous other evergreen forest angiosperm genera e.g. *Vepris* Comm. ex A.Juss. (Rutaceae, Cheek & Onana 2021), *Saxicolella* Engl. (Podostemaceae, Cheek et al. 2022), *Cola* Schott & Endl. (Malvaceae-Sterculioideae, Cheek et al. 2020a) and *Uvariopsis* Engl. & Diels (Annonaceae, Couvreur et al. 2022; Gosline et al. 2022). The Interval has the highest species and generic diversity of flowering plants per degree square in tropical Africa (Barthlott et al. 1996; Dagallier et al. 2020) including endemic genera such as *Medusandra* Brenan (formerly Medusandraceae, now Peridiscaceae, Soltis et al. 2007; Breteler et al. 2015), and new genera to science are still being discovered e.g. *Korupodendron* Litt & Cheek (Vochysiaceae, Litt & Cheek 2002) and *Kupea* Cheek & S.A.Williams (Triuridaceae, Cheek et al. 2003).

The species of *Memecylon* in Guineo-Congolian Africa (sensu White 1983) are classified as belonging to subg. *Mouririoidea* (Jacq.-Fél.) R.D. Stone or subg. *Memecylon* with the latter further subdivided into six endemic sections: *Afzeliana* Jacq.-Fél., *Diluviana* R.D. Stone, *Felixiocylon* R.D. Stone, *Germainiocylon* R.D. Stone, *Polyanthema* Engl. s.str., and *Sitacylon* R.D. Stone (Stone 2014a). Section *Afzeliana* is distinguished by its ellipsoid to oblong fruits that are often white when young, turning blue or purple at maturity; the remaining sections have globose fruits that are usually green when immature. The section *Afzeliana* was recently revised, recognising 20 species, 17 of which are in Cameroon (Stone et al. 2008), and *Memecylon emancipatum* R.D. Stone from Liberia has since been recognised (Stone 2014b). *Afzeliana* is also a monophyletic group in phylogenetic analyses of DNA sequences (Stone 2014a).

In this paper, we formally describe *Memecylon ebo*, a further new species of sect. *Afzeliana* that is narrowly endemic within the Cross-Sanaga River Interval, specifically the Ebo forest in Littoral Region, Cameroon. Several photos taken in the Ebo Forest of an unusual yellow-petalled Melastomataceae-Olisbeoideae with jade-green stamens, subtrinervate leaf-blades, subsessile inflorescences and highly contracted pedicels (Fig. 1) were found by MC in early 2023 when reviewing photographic images for use in the book Important Plant Areas of Cameroon (Murphy et al. 2023). These photos were referred to RDS who identified them as a new species of *Memecylon* sect. *Afzeliana* (Stone et al. 2008). The specimen they were linked to, *Prenner* 23 (K, YA), had erroneously been identified as *Memecylon zenkeri* Engl. which has long inflorescences bearing white petalled, blue-staminate flowers on long pedicels (Fig. 2). However, the two species are vegetatively highly similar, both having quadrate-alate young internodes, thickly leathery leaves with conspicuous transverse veins united with a pair of looping intramarginal nerves. Searches of all specimens of *Memecylon zenkeri* at K and of 173 specimen images on GBIF (gbif.org) brought to light six other flowering and fruiting specimens that matched *Prenner* 23, all from a small area in the NE quadrant of the Ebo Forest. The ellipsoid fruits of the specimens matching *Prenner* 23 confirmed placement in *Memecylon* sect. *Afzeliana*.

**Fig. 1.**
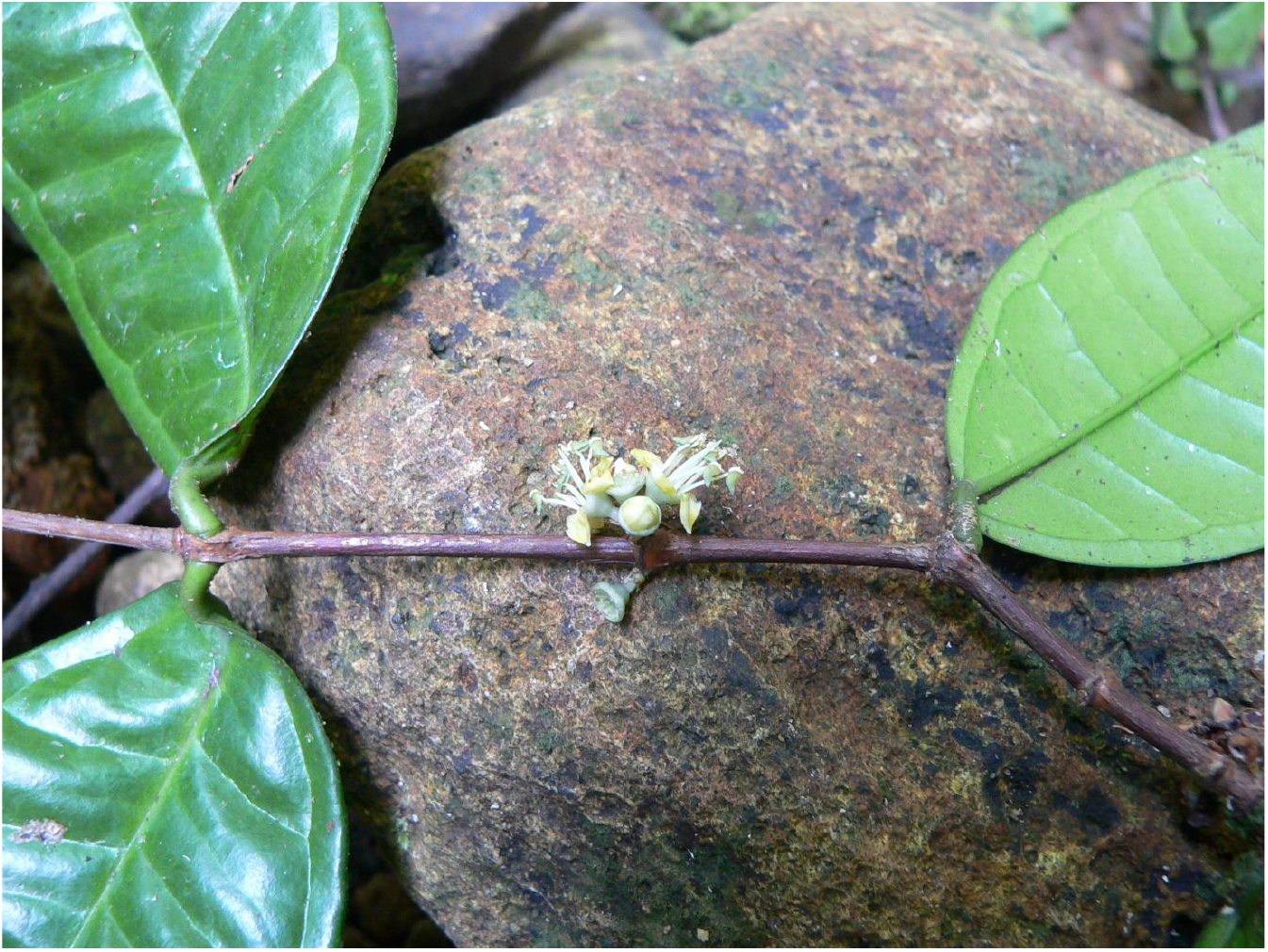
*Memecylon ebo* Flowering branch of *Prenner* 23. Photo by Gerhard Prenner.

**Fig. 2.**
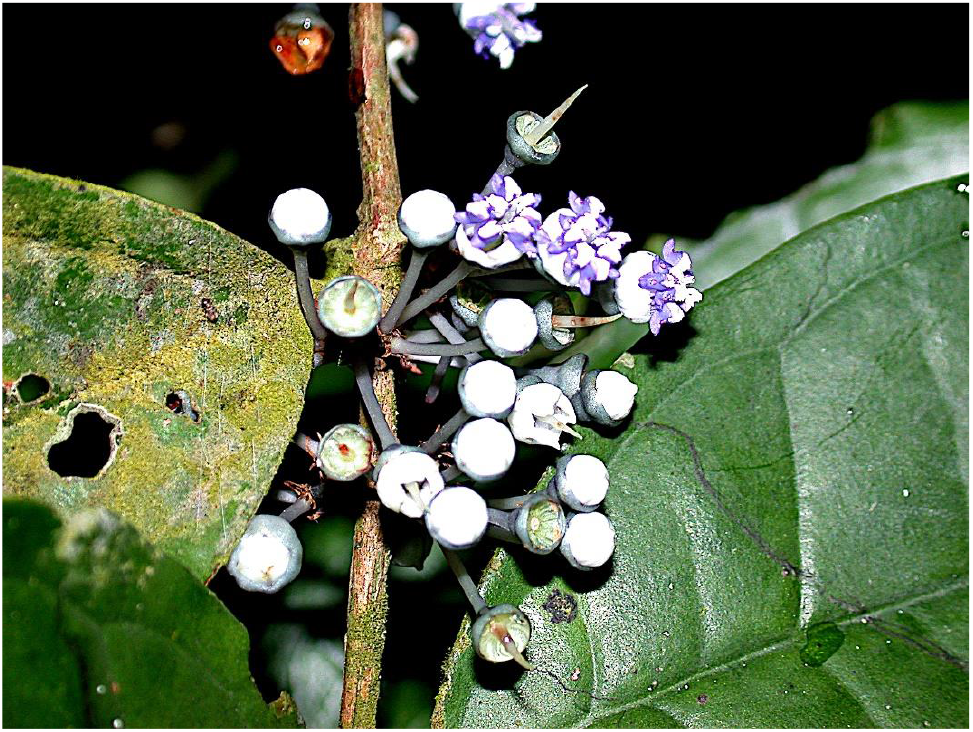
*Memecylon zenkeri* Flowering branch. Note the “star flower” floral architecture and the blue anther connectives with oil glands, all characteristic of *Memecylon* sect. *Afzeliana* (Stone et al. 2008). Photo by R.D. Stone

We also review and update information on the endemic plant species of Ebo Forest, recently designated as an Important Plant Area of Cameroon (Murphy et al. 2023) and under immense threat of commercial logging.

## Materials & Methods

This study is based on herbarium specimens. All specimens cited have been seen by us unless indicated as “n.v.”. The methodology for the surveys in which most of the specimens were collected is given in Cheek & Cable (1997). Herbarium citations follow Index Herbariorum (Thiers *et al*. continuously updated), nomenclature follows Turland *et al*. (2018) and binomial authorities follow IPNI (continuously updated). The Flore du Cameroun volume for Melastomataceae (Jacques-Félix 1983) followed by the taxonomic treatment of *Memecylon* sect. *Afzeliana* (Stone et al. 2008) were the principal reference works used to determine the identifications of the specimens of what proved to be the new species. Material of the suspected new species was identified by comparing morphologically with protologues, reference herbarium specimens (Cheek in Davies et al. 2023), including type material of *Memecylon* sect. *Afzeliana* principally at K, but also using material and online images from BR, MO, P and YA, including all 173 images on gbif.org (accessed 15 April 2023) of *Memecylon zenkeri*. The description was made following the terms used in Beentje & Cheek (2003) and the format of Stone et al. (2008) and Stone & Cheek (2018).

The conservation assessment was made in accordance with the categories and criteria of IUCN (2012). Herbarium material was examined with a Leica Wild M8 dissecting binocular microscope fitted with an eyepiece graticule measuring in units of 0.025 mm at maximum magnification. The drawing was made with the same equipment using a Leica 308700 camera lucida attachment.

### Taxonomic treatment

The new species from Ebo represented by *Prenner* 23, because it has quadrate-alate stems in which the nodes are <2 x the diameter of the internodes, and leaves which are both subtrinervate and with a pair of looping intramarginal nerves connecting with the transverse veins which are impressed above and prominent below, keys out to *Memecylon zenkeri* at couplet 15 in the revision of Stone et al. (2008). Characters separating the two species are indicated in table 1 below.

**Table 1.**
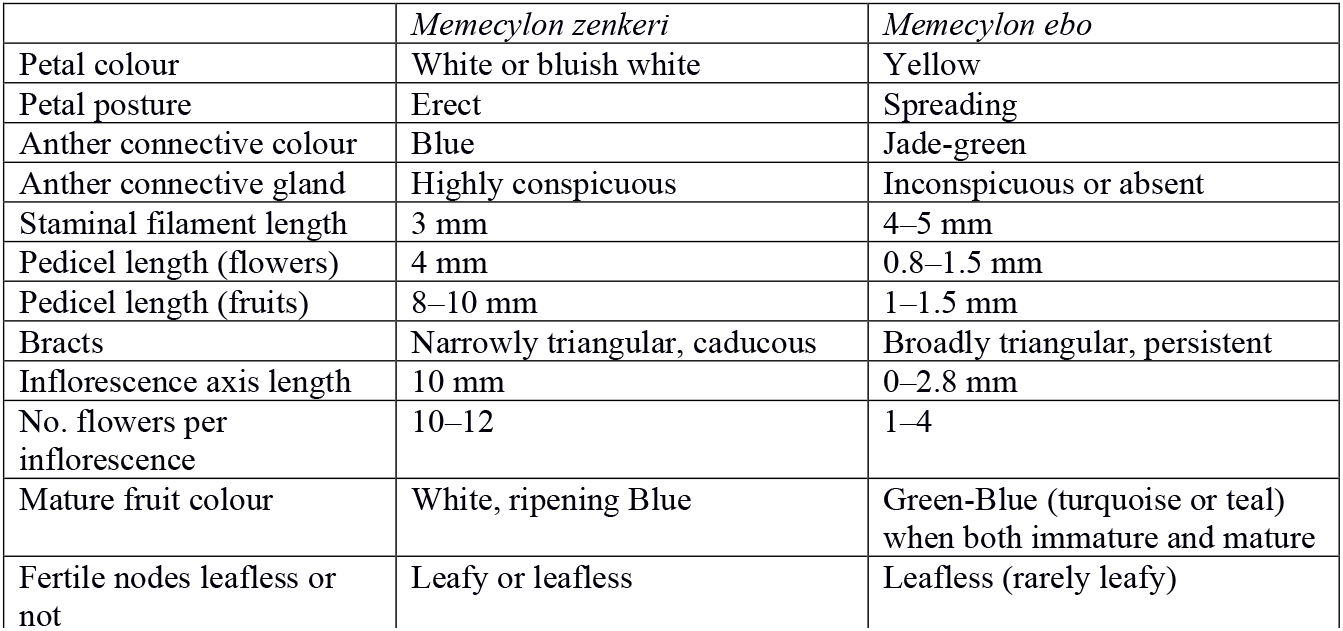
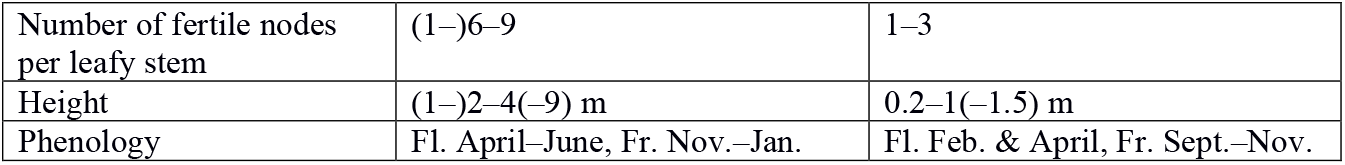
Diagnostic differences separating *Memecylon zenkeri* from *M. ebo*. Data for *M. zenkeri* from *Fl. Cameroun* (Jacques-Félix 1983) and from specimens at K (*n*=40).

|  | *Memecylon zenkeri* | *Memecylon ebo* |
| --- | --- | --- |
| Petal colour | White or bluish white | Yellow |
| Petal posture | Erect | Spreading |
| Anther connective colour | Blue | Jade-green |
| Anther connective gland | Highly conspicuous | Inconspicuous or absent |
| Staminal filament length | 3 mm | 4–5 mm |
| Pedicel length (flowers) | 4 mm | 0.8–1.5 mm |
| Pedicel length (fruits) | 8–10 mm | 1–1.5 mm |
| Bracts | Narrowly triangular, caducous | Broadly triangular, persistent |
| Inflorescence axis length | 10 mm | 0–2.8 mm |
| No. flowers per inflorescence | 10–12 | 1–4 |
| Mature fruit colour | White, ripening Blue | Green-Blue (turquoise or teal) when both immature and mature |
| Fertile nodes leafless or not | Leafy or leafless | Leafless (rarely leafy) |
| Number of fertile nodes per leafy stem | (1–)6–9 | 1–3 |
| Height | (1–)2–4(–9) m | 0.2–1(–1.5) m |
| Phenology | Fl. April–June, Fr. Nov.–Jan. | Fl. Feb. & April, Fr. Sept.–Nov. |

**Memecylon ebo** *R. D. Stone & Cheek*, **sp. nov**. Typus: Cameroon, Littoral Region, Yingui, Bekob village, Yabassi, 4°25’ 14”N, 10°24’ 45.0”E, alt. 1040 m, forest understory, fls, 28 Apr. 2005, *Tchiengue* 2107 with Ekwoge, Horwath, Hoffmann (holotypus K barcode K001243169; isotypus YA).

#### Evergreen understorey shrub

0.2–1(–1.5) m tall, entirely glabrous. Branchlets slender, the youngest 1–3 internodes quadrangular in cross section and narrowly alate (wings c. 0.20–0.25 mm wide) becoming terete with age, epidermis subglossy, midbrown; internodes (1.3–)1.5–2.8(–3.1) cm long, 1–2 mm diam., nodes swollen 1.8–3.5 mm diam. *Leaves* on petioles 2–4(–6) x 1 mm, the petioles twisted, channelled on the adaxial side. Blades thickly coriaceous, dark green and shining on the upper surface, paler beneath, drying black above and midbrown beneath, lanceolate-elliptic or narrowly elliptic-oblong in outline, (8.4–)10–14.5(–15) x (2.7–)3.3–5(–5.5) cm, base acute to broadly acute, acuminate at apex, the acumen narrowly triangular (0.8–)1–1.5 x 0.2–0.3 cm; midnerve, lateral and transverse nerves strongly impressed on the upper surface, sub-bullate, prominent on the lower, the lateral nerves diverging from the midnerve at the base of the blade, forming arches between the junctions with the transverse veins, 2–5 mm from the margin; transverse veins 9–16 pairs, of about the same thickness as the laterals, prominent on the lower surface, network of smaller venules absent or inconspicuous. *Inflorescences* subsessile cymules, 1– 4-flowered, 0.5–0.6 cm long (including flowers), borne mostly at the leafless, slightly thickened nodes of otherwise leafy branches (rarely at the leafy nodes), 1–7 nodes distant from the stem apex, 1–3 flowering/fruiting nodes per leafy stem. Inflorescence axis 0–2.8 x 1.5 mm, unbranched; bracts opposite, in pairs, with up to 3 pairs of bracts per axis, spreading, triangular convex, 1.2–1.5 x 1.3 mm, apex often with a mucro, bases often united, sheathing the axis, forming a boat shaped structure, persistent in fruit (Fig 3E). *Flowers* subsessile, pedicels partly concealed by the subtending bracts, 0.8–1.5 x 0.8 mm; hypantho-calyx pale greenish white, obconic, c. 2 mm long x 3.5 mm diam., the calyx margin entire, lacking lobes, the inner surface with eight radial ridges in pairs, flanking the bases of the staminal filaments (Fig.3J). Petals pale yellow (Fig. 1), 4, spreading to slightly recurved, outer petals subtriangular, 2.5–3 x 3.5 mm, apex obtuse, base shortly and broadly unguiculate, margins involute; inner petals as the outer but smaller, quadrangular, 2–2.1 mm wide, apex acute. Stamens long-exserted on slender white cylindrical filaments 4–5 x 0.2–0.25 mm; anther cells white, oblong, (0.6–)1.5 x 0.6 mm, the connective jade green, extending dorsally, conical in dorsal view, excavated ventrally (1.5–) 2 x (0.4–)0.7–1 mm, in side view in life slightly hook shaped (Fig. 3J) and outwardly curved, lacking a dorsal keel, the oil-gland absent or diffuse and very inconspicuous appearing as a dark discolouration between the midpoint and the extremity of the connective. Style slender, 5–7 x 0.6–0.7 mm, tapering to an acute apex.

**Fig. 3.**
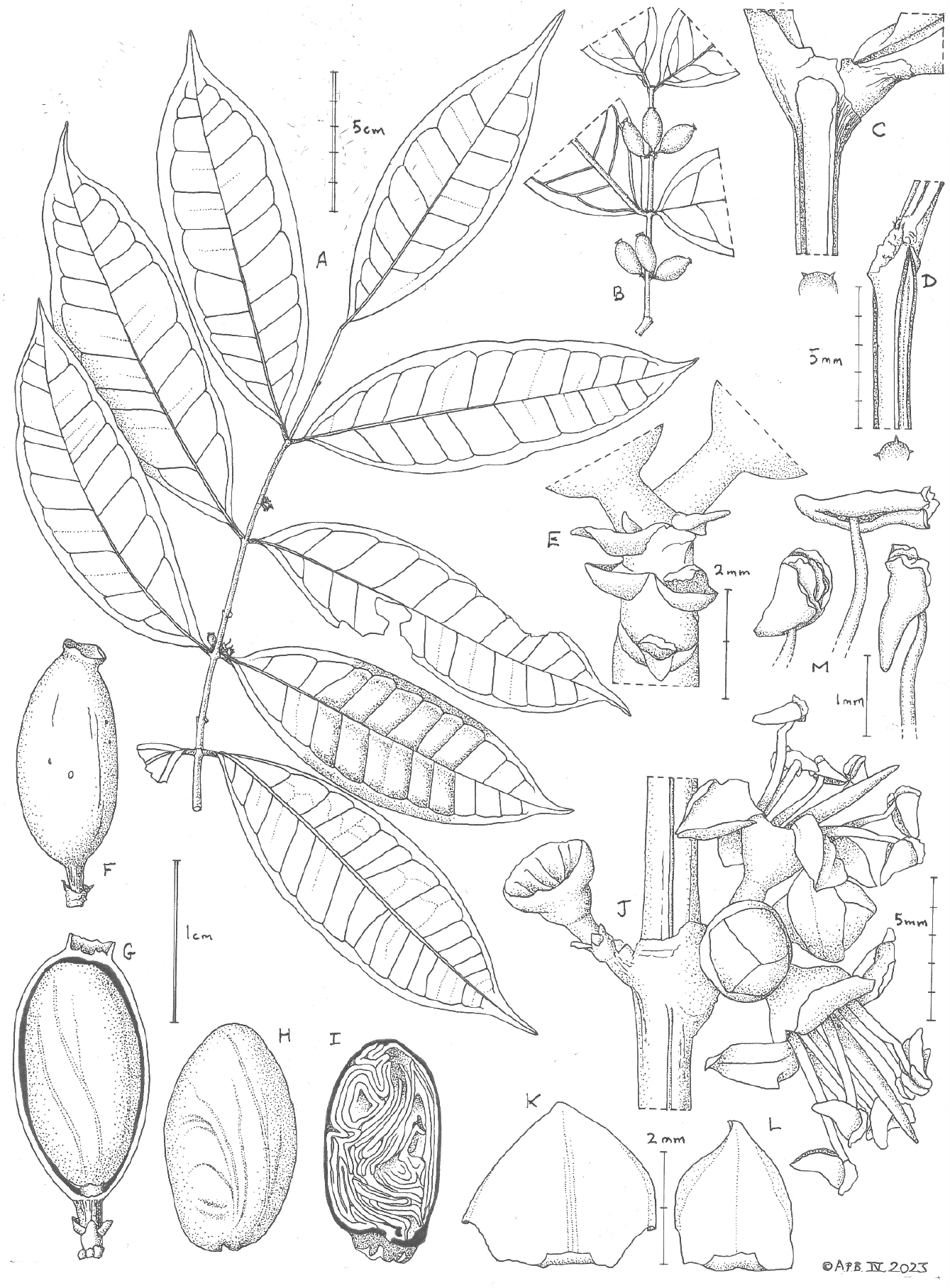
*Memecylon ebo* **A** habit, flowering branch; **B** habit, flowering branch; **C & D** detail of stem apex, leaf bases and internode, also transverse sections of stem showing four wings; **E** inflorescence axis, showing bract pairs, pedicels with base of two fruits; **F** fruit, showing sheathing bract pairs; **G** fruit, rehydrated, with seed exposed; **H** rehydrated seed; **I** seed in longitudinal section showing the folded cotyledons; **J** two inflorescences at leafless node; **K** outer petal, outer (abaxial) surface; **L** inner petal, inner (adaxial surface); **M** three rehydrated anthers. **A & K**-**M** from *Tchiengue* 2107; **B**-**D & F** from *Xanthos* 297; **E, G**-**I** from *Osborne* 198; **J** from *Prenner* 23. All drawn by ANDREW BROWN

*Fruit* white when young, blue-green at maturity, ellipsoid-oblong (9–)12–16 x (6–)7–9 mm long, crowned by the persistent but non-accrescent calyx, calyx broadly funnel-shaped c. 1 x 3 mm; mesocarp fleshy-leathery, seedcoat grey brown, smooth, slightly smaller than the fruit (Fig. 3H), embryo with cotyledons foliaceous, highly folded (Fig. 3I).

##### RECOGNITION

[diagnosis] *Memecylon ebo* differs from all other species of *Memecylon* in the yellow petals and jade green anther-connectives.

In the absence of flowers, *Memecylon ebo* has been confused with the sympatric *M. zenkeri* Engl., also of sect. *Afzeliana*. However, in fruit the first species can be recognised by the infructescence axes being subsessile or very short, 0–2.8 mm long (vs c.10 mm long), and the fruiting pedicels very short, 1–1.5 mm (vs 8–10 mm long),

##### DISTRIBUTION

Cameroon. Endemic on current evidence to the NE quadrant of Ebo Forest, Littoral Region.

##### ADDITIONAL COLLECTIONS. CAMEROON. Littoral Region

Yabassi, Yingui, Ebo Forest, Bekob village, Ndokbaembi on N-S ridge, along century old abandoned German rd, fr. 21 April 2005, *Cheek* 12435 with Morgan, Corcoran, Jonas (CAS, K000460187, SCA n.v., YA n.v.); ibid., towards the river, Nolokbayembi path, fl. 17 Feb. 2006, *Prenner* 23 with Fenton, Alobuede (K001243168, YA n.v.); ibid. off S transect along river, fr. 15 Sept. 2006, *Osborne* 62 with Engomgwie, Mam, Morgan (K000341249, YA); ibid., Bekob drinking stream trail, 1550 m along, fr. 27 Oct. 2006, *Osborne* with Beheng 198 (K000341246, YA n.v.); ibid., Ndogbayembe trail 1550 m from Bekob camp, fr. 11 June 2008, *MacKinnon* 174 (K001243166, MO n.v., YA n.v.); ibid., Bekob, 525 m along Decamb trail, fr. 30 Nov. 2013, *Xanthos* 297 with van der Burgt, Ngansop, Tchiengue (G n.v., K 001286641, YA n.v.).

##### HABITAT & ECOLOGY

Intact, undisturbed, submontane evergreen (cloud) forest; 710–1040 m alt.

##### CONSERVATION STATUS

*Memecylon ebo* is assessed here as Critically Endangered CR B1ab(iii) +B2ab(iii) using the IUCN (2012) criteria. This is because it is known from a single threat-defined location with area of occupancy and extent of occurrence both of about 4 km^2^ in the NE quadrant of the Ebo forest around the Banen village of Bekob, currently uninhabited. This is one of the most intensively surveyed parts of the Ebo forest because Bekob was for many years the main research centre for biologists in the forest. Within this small area it appears not infrequent nor inconspicuous since seven collections were made over nine years (2005–2013) by six collectors (see above). Many memecyloid species are both restricted in their range and infrequent within it (Cheek pp. 212–222 in: Onana & Cheek 2011), and this is supposed by us to be the case with *Memecylon ebo*. Surveys elsewhere in Ebo, for example around Njuma in the NW quadrant of Ebo Forest, also intensively surveyed, have failed to find this species.

It is possible that *Memecylon ebo* will yet be found at additional locations in Cameroon outside Ebo. However, while surveys have not been exhaustive, many thousands of specimens have been collected in areas to the north, south, west and east of Ebo (Cheek et al. 1992; Cheek *et al*. 1996; Cable & Cheek 1998; Cheek *et al*. 2000; Maisels *et al*. 2000; Chapman & Chapman 2001; Harvey *et al*. 2004; Cheek *et al*. 2004; Cheek *et al*. 2006; Cheek *et al*. 2010; Harvey *et al*. 2010; Cheek *et al*. 2011). Given the vegetative similarity of *Memecylon ebo* to *M. zenkeri*, all 173 specimen images of that species on gbif.org were checked in case further specimens of the new species might be found under this name, but no new additional records were detected. In flower or fruit the two can be distinguished easily by the long pedicels of the second species, versus subsessile or very short in the first species.

If *Memecylon ebo* occurs elsewhere, that is most likely to be in the Bakossi area to the west of Ebo in SW Region, especially Mt Kupe to the NW since several threatened range-restricted species are confined to these two areas, e.g. *Costus kupensis* Maas & H. Maas (Maas-van der Kamer et al. 2016), *Coffea montekupensis* Stoff. (Stoffelen *et al*. 1997), *Microcos magnifica* Cheek (Cheek 2017a), *Dovyalis* sp. 1 of Mt Kupe (Cheek et al. 2004), *Impatiens frithii* Cheek (Cheek & Csiba 2002). However, since Mt Kupe has been intensively collected, we consider this unlikely.

The Ebo forest, while recently designated as an Important Plant Area (Murphy et al. 2023) is not protected and a large part was designated as a logging concession in Feb. 2021 (Lovell 2020). Although this was suspended by the President of Cameroon in Aug. 2021 (Kew Science News 2020), the forest is now immediately threatened by a further logging concession in 2023 where logging has started at the eastern and southern edges.

Additional threats are clearance of forest habitat for oil palm plantations and for an open cast iron-ore mine (Cheek *et al*. 2018a).

##### PHENOLOGY

Flowering in Feb. and April. Fruits ripening in Sept.–Nov.

##### ETYMOLOGY

Named for the forest of Ebo in Littoral Region, Cameroon, to which *Memecylon ebo* appears endemic.

##### VERNACULAR NAMES & USES

None are recorded.

##### NOTES

*Memecylon ebo* appears to be relatively frequent around the Bekob village (abandoned) former research base where all seven known specimens occur in 4–8 square kilometres. In contrast, the vegetatively similar *M. zenkeri* is much less frequent but more widespread, with eight specimens recorded scattered mostly thinly over the several hundred square kilometres within the Ebo forest that have been surveyed, with a cluster of three specimens near the former Juma camp, in the NW quadrant of Ebo. One specimen of *M. zenkeri* (*Lockwood* 44), is recorded near Bekob, so that the two species appear to be sympatric. *Memecylon zenkeri* is a relatively widespread species, occurring from SE Nigeria to Gabon.

Apart from these two species, only one other species of the genus, *Memecylon viresecens* Hook. f., has been recorded from Ebo thus far, with a single record sympatric with *M. ebo* near Bekob, *MacKinnon* 213 (K, YA).

That the connective oil-gland is absent or diffuse and very inconspicuous in *Memecyon ebo* is unusual in the genus. However, not all members of sect. *Afzeliana* have the oil-gland. It is also reduced or absent in *M. mamfeanum* (Jacq.-Fél.) R.D.Stone, Ghogue & Cheek, M. hyleastrum R.D.Stone & Ghogue, *M. bakossiense* R.D.Stone, Ghogue & Cheek, and *M. rheophyticum* R.D.Stone, Ghogue & Cheek.

### The endemic plant species of Ebo Forest, Littoral Region

The Ebo Forest, a former proposed National Park, covers c. 1,400 km^2^ of lowland and submontane forest, including important inselberg and waterfall areas, with an altitudinal range of 130–1115 m alt. and a rainfall of 2.3 – 3.1 m per annum (Abwe & Morgan 2008; Cheek et al. 2018a). To date over 100 globally threatened plant species have been documented including 23 new to science (Murphy et al. 2023), of which *Memecylon ebo* is the tenth that is globally endemic to Ebo (see Table 2).

**Table 2.** The strictly endemic plant species of Ebo, and those near-endemic, i.e. also at one or two other locations.

| Species name<br>(family) | Strictly<br>Endemic<br>to Ebo | Also at (location<br>1) | Also at<br>(location 2) | IUCN<br>Conservation<br>Assessment if<br>available on<br>iucnredlist.org | Reference<br>(* = newly<br>recorded at<br>Ebo in this<br>paper) |
| --- | --- | --- | --- | --- | --- |
| *Afrothismia*<br>*fungiformis*<br>(Afrothismiaceae) |  | Mt Kupe |  | EN | Cheek *et al.* (2023a) |
| *Piptostigma*<br>*submontanum*<br>(Annonaceae) |  | Rumpi Hills | Bakossi Mts | EN | Ghogue *et al.* (2017) |
| *Uvariopsis*<br>*dicaprio*<br>(Annonaceae) | X |  |  | CR | Gosline *et al.* (2022) |
| *Crateranthus cameroonensis*<br>(Lecythidaceae) | X |  |  | CR | Prance & Jongkind (2015) |
| *Inversodicraea ebo*<br>(Podostemaceae) | X |  |  | CR | Cheek et al. (2017) |
| *Costus kupensis*<br>(Costaceae) |  | Mt Kupe |  | EN | Maas-van der Kamer et al. (2016) |
| *Microcos magnifica*<br>(Grewiaceae) |  | Mt Kupe |  | EN | Cheek (2017a) |
| *Pseudohydrosme ebo* (Araceae) | X |  |  | CR | Cheek et al. (2021) |
| *Palisota ebo*<br>(Commelinaceae) | X |  |  | CR | Cheek et al. (2018a) |
| *Impatiens frithii*<br>(Balsaminaceae) |  | Bakossi Mts | Mt Etinde-Mt Cameroon | EN | Cheek & Csiba (2002)* |
| *Impatiens banen*<br>(Balsaminaceae) | X |  |  | n/a | Cheek et al. (2023b) |
| *Kupeantha ebo*<br>(Rubiaceae) | X |  |  | CR | Cheek et al. (2018b) |
| *Ardisia ebo*<br>(Primulaceae) | X |  |  | CR | Cheek & Xanthos (2012) |
| *Psychotria pachycalyx*<br>(Rubiaceae) |  | Rumpi Hills | Bata, Rio Muni | n/a | Lachenaud (2019) * |
| *Berlinia korupensis*<br>(Leguminosae) |  | Korup |  | CR | Mackinder & Pennington (2011)* |
| *Talbotiella ebo*<br>(Leguminosae) |  | Loum Forest Reserve | Mone Forest Reserve | EN | Mackinder et al. (2010) |
| *Dichapetalum korupinum*<br>(Dichapetalaceae) |  | Korup |  | CR | Breteler (1996) * |
| *Phyllanthus caesiifolius*<br>(Phyllanthaceae) |  | Bakossi |  | CR | Hoffmann & Cheek (2003) * |
| *Phyllanthus nyale*<br>(Phyllanthaceae) |  | Bakossi |  | CR | Hoffmann & Cheek (2003) * |
| *Rhaptopetalum breterlei*<br>(Lecythidaceae) |  | Nguélémendouka |  | CR | Prance & Jongkind (2015) * |
| *Aulacocalyx camerooniana*<br>(Rubiaceae) |  | Akom II |  | CR | Sonké et al. (2005); Cheek (2017b)* |
| *Coffea leonimontana*<br>(Rubiaceae) |  | S of Kompina<br>(Littoral) where<br>likely extinct |  | CR | Stoffelen<br>et al.<br>(1996)* |
| *Vitex yaundensis*<br>(Lamiaceae) |  | Yaoundé | Mungo<br>River Forest<br>Reserve | CR | Pollard<br>(2004)* |
| *Memecylon ebo*<br>(Melastomataceae) | X |  |  | n/a | This paper |

Ebo was recently evidenced (Kew Science News, 2020) as an Important Plant Area (IPA). Of the 49 IPAs in Cameroon, Ebo has the highest number (23) of documented Critically Endangered (CR) plant species (IUCN global assessments), i.e. those with the highest level of global threat and closest to extinction, after Ngovayang (Bipindi) which has 24 (Murphy et al. 2023: 23, table 3). With the addition of *Memecylon ebo*, here assessed as CR also, Ebo will equal Ngovayang, and with the description and assessment of additional narrow endemics in the course of preparation for publication including in the genera *Ardisia* Sw., *Begonia* L., *Chassalia* Comm. ex Poir., *Cola, Microcos* L., *Mitriostigma* Hochst., it will soon have the highest number of CR species in Cameroon, and possibly in all of tropical Africa. Most of Ebo’s CR species are endemic to Ebo (see Table 2 detailing the 25 endemic or near endemic plant species of Ebo). However, several other CR species of Ebo occur at more than two other additional sites (see Murphy et al. 2023; 169 – 177) e.g. *Belonophora ongensis* S.E.Dawson & Cheek (Rubiaceae, Cheek & Dawson (2000)). Other point endemic CR species occur just outside the boundary of Ebo, e.g. *Kupeantha yabassi* M.G.Alvarez & Cheek (Rubiaceae, Alvarez-Aguirre et al. 2021). Ebo also holds the only Cameroonian location for globally threatened species such as *Nothodissotis barteri* (Hook.f.) Ver.-Lib. & G.Kadereit (Veranso-Libalah et al. 2019).

## Discussion

The publication in this paper of a further endemic species of plant for the Ebo forest emphasises further its global importance for conservation. No other site in Cameroon has more Critically Endangered plant species now than Ebo.

Cameroon has the highest number of globally extinct plant species of all countries in continental tropical Africa (Humphreys et al. 2019). The extinction of species such as *Oxygyne triandra* Schltr. (Thismiaceae, Cheek et al. 2018b) and *Afrothisia pachyantha* Schltr. (Afrothismiaceae, Cheek & Williams 1999; Cheek et al. 2019a; Cheek et al. 2023a) are well known examples, recently joined by species such as *Vepris bali* Cheek (Rutaceae, Cheek et al. 2018c), *Vepris montisbambutensis* Onana (Onana & Chevillotte 2015) and *Ardisia schlechteri* Gilg (Murphy et al. 2023). However, another 127 potentially globally extinct Cameroon species are documented (Murphy et al. 2023: 18 – 22).

It is critical now to detect, delimit and formally name species as new to science, since until they are scientifically recognised, they are essentially invisible to science, and only when they have a scientific name can their inclusion on the IUCN Red List be facilitated (Cheek et al. 2020b). Most (77%) species named as new to science in 2023 are already threatened with extinction (Brown et al. 2023). Many new species to science have evaded detection until today because they have minute ranges which have remained unsurveyed until recently, as was the case with *Memecylon ebo*. However, there are exceptions (Cheek & Etuge 2009; Cheek et al. 2019b).

If further global extinction of plant species is to be avoided, effective conservation prioritisation, backed up by investment in protection of habitat, ideally through reinforcement and support for local communities who often effectively own and manage the areas concerned, is crucial. Important Plant Areas (IPAs) programmes, often known in the tropics as TIPAs (Darbyshire et al. 2017; Murphy et al. 2023) offer the means to prioritise areas for conservation based on the inclusion of highly threatened plant species, among other criteria. Such measures are vital if further species extinctions are to be avoided of narrowly endemic, highly localised species such as *Memecylon ebo*.

## Supporting information

Supplemental file Holotype image Memecylon ebo

## Acknowledgements

The authors are especially grateful to Dr Gerhard Prenner for providing the photos that allowed the detection of the new species published in this paper and for permitting their reproduction.

This paper was completed as part of the Cameroon TIPAs (Tropical Important Plant Areas) project at RBG, Kew, which is supported by Players of Peoples Postcode Lottery. We thank Lydia Burns and Penny Appelbe of Kew’s Foundation for making this possible. The second author’s contribution to this paper was made possible by visits from Cameroon to RBG, Kew, U.K. sponsored by the Bentham-Moxon Trust of RBG, Kew. Many of the specimens of the new species were collected by dedicated independent botanists from the UK supporting the inventory of Ebo: Martin Xanthos, Jo Osborne, Lorna Mackinnon, Helen Lockwood and Emma Fenton.

We thank the Bentham Moxon Trust, the Darwin Initiative, and Earthwatch Europe for supporting the surveys by sponsoring the travel and accommodation costs.

This paper is a result of the partnership between RBG, Kew and MINRESI-IRAD-National Herbarium of Cameroon, Yaoundé and a series of Memoranda of Collaboration that began in 1996. We thank the late Dr Benoît Satabié, Drs Gaston Achoundong, Florence Ngo Ngwe, Eric Nana, Jean Lagarde Betti, the current and former directors or acting Directors, of IRAD-National Herbarium of Cameroon, Yaoundé, for expediting the collaboration between our two institutes.

We further thank the National Herbarium of Cameroon-IRAD-for permission to collect specimens in Cameroon, to the Ebo Forest Research Project supported by the San Diego Zoo Wildlife Alliance (SDZWA) and the Banen people for supporting our botanical surveys in the Ebo forest Cameroon.

Two anonymous reviewers are thanked for comments on an earlier version of the manuscript.

The authors declare no conflicts of interest.

