## Supplementary figures and images for "The endemic plant species of Ebo Forest, Littoral Region, Cameroon with a new Critically Endangered cloud forest shrub, *Memecylon ebo* (Melastomataceae-Olisbeoideae)"

### Supplemental file Holotype image Memecylon ebo

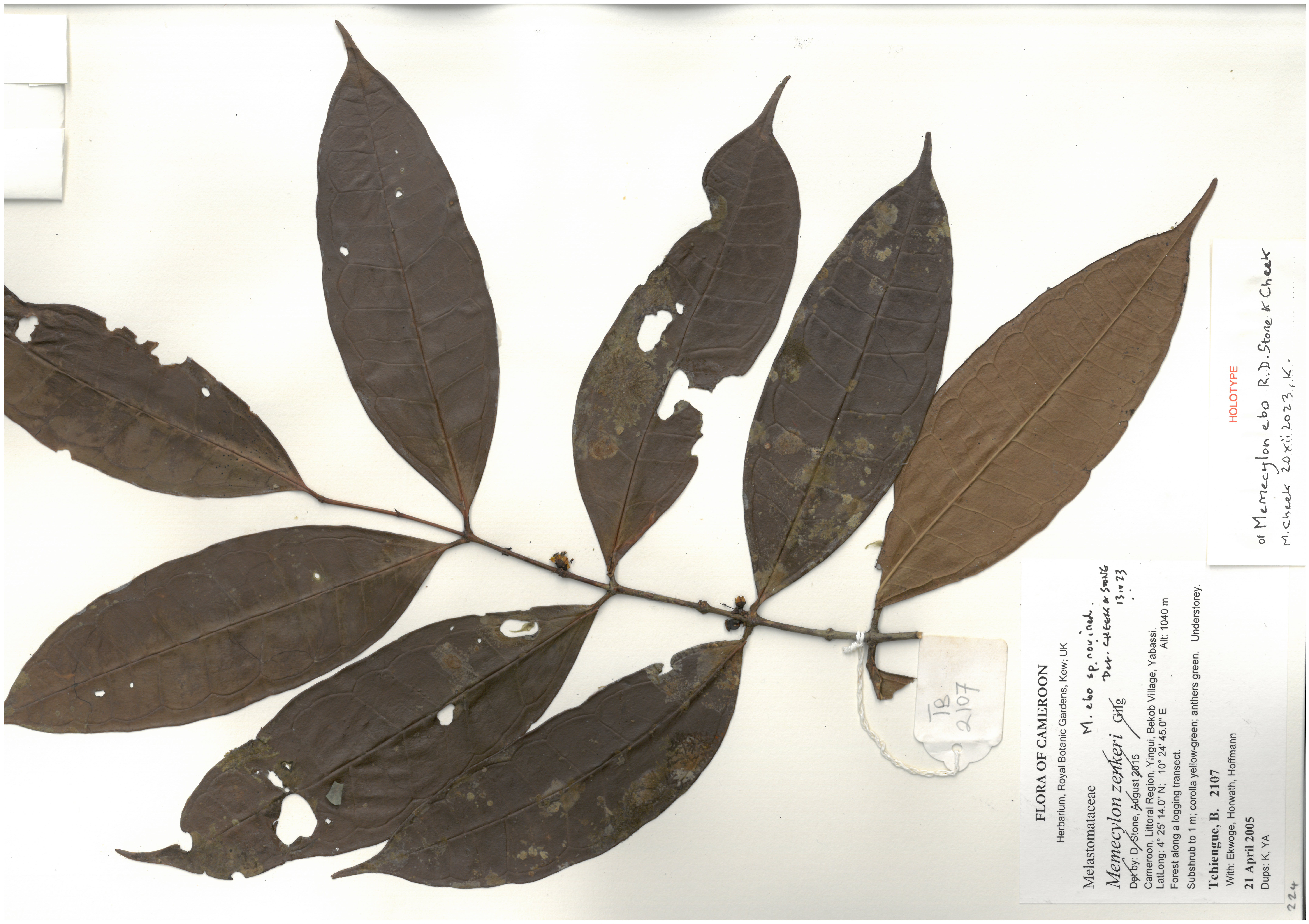
